# Extracellular ssDNA from *Pittosporum tobira* Exerts Strong Insecticidal Activity on *Coccus hesperidum*: a natural parallel to ‘genetic zipper’ technology

**DOI:** 10.1101/2025.11.18.689116

**Authors:** Vol Oberemok, Kate Laikova, Nikita Gal’chinsky, Jamin Ali, Natalia Petrishina, Yekaterina Yatskova, Ilyas Chachoua

## Abstract

Beyond its function as a carrier of hereditary information, recent research has uncovered novel properties of extracellular DNA, including its role in the adaptation to the environment when released from plants. The secreted DNA has been shown to exert insecticidal effects against insect pests, which play an adaptive role in plant-insect interactions, particularly in regulating populations of economically important sap-feeding insects. The molecular mechanisms underlying this insecticidal effect are underinvestigated and remain largely unknown. Therefore, there is a need for more efforts to uncover these mechanisms to better understand the plant-pest interactions, which would provide new insights into natural pest control strategies and inspire biotechnological applications. In the current study, we show that *Pittosporum tobira* (*P. tobira*) secretes single-stranded DNA (ssDNA) that exerts an insecticidal effect on *Coccus hesperidum* (*C. hesperidum*). We collected extracellular DNA from *P. tobira* leaves and tested its potential insecticidal effect by applying it to *C. hesperidum*, which is a well-known pest that causes damage to *P. tobira*. Our results revealed that the outermost layer of the leaf cuticle of *P. tobira* predominantly contains ssDNA of approximately 100 nt in length, originating from both chloroplast and nuclear genomes. This DNA exhibited pronounced insecticidal activity against *C. hesperidum*, with chloroplast-derived sequences significantly enriched compared to the total DNA in intact plant cells. These findings suggest that the microevolution of the *P. tobira* nucleome and plastome contributed to the formation of extracellular DNA with insecticidal properties (eci-DNA), which is part of its defence strategy against insect pests. Moreover, in this article for the first time we show that antisense DNA (oligonucleotide insecticide Coccus-11) is capable of activating insect retrotransposons and upregulating their RT-RNase H, a crucial enzyme for the DNA containment mechanism and successful action of oligonucleotide insecticides. Notably, the laboratory-developed ssDNA-based ‘genetic zipper’ technology, designed for sustainable pest management, possesses characteristics similar to eci-DNA found in nature, highlighting a potential natural parallel to this biotechnological approach for sustainable pest management.

**Graphical Abstract:** 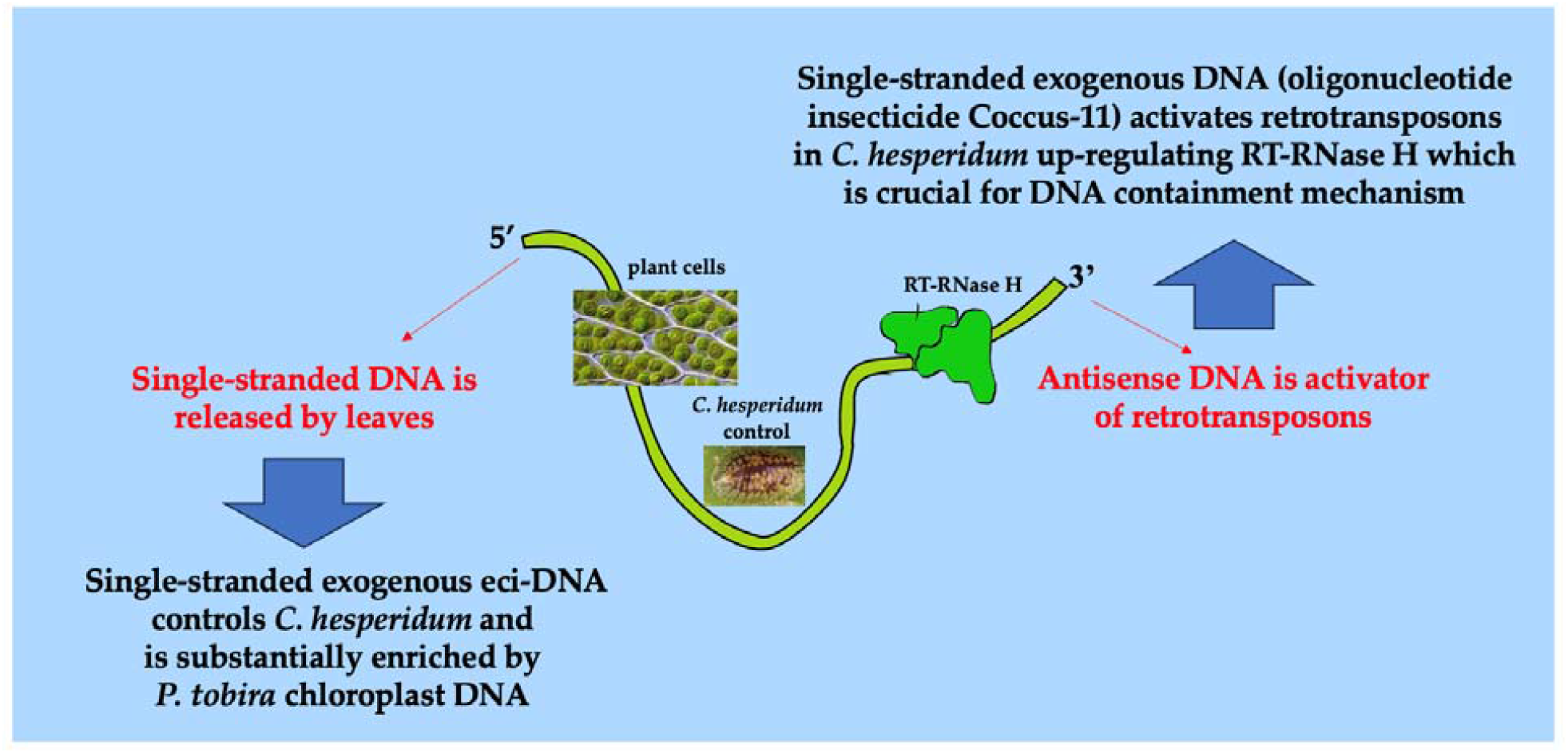

## 1. Introduction

Recent advancements in molecular biology techniques have shed light on the structure and functions of biomolecules such as DNA, RNA and proteins. DNA has recently been shown to be an active tool for communication and interaction between cells and different organisms, including plants and insects. We are gradually beginning to understand the importance of DNA as a programmable molecule for the interactions of organisms with each other, the functioning of distinct ecosystems, and the maintainance of the balance of the biosphere as a whole. Plants, as the main primary producers, use their extracellular DNA (ecDNA) as a versatile protective macromolecule to counteract environmental factors. Although it is not entirely clear how exactly ecDNA is formed in cells, its secretion mainly results from either cell lysis or active release. Secreted ecDNA exists in more or less fragmented forms and can be found as both single-stranded DNA (ssDNA) and double-stranded DNA (dsDNA) forms. Current knowledge suggests that the formation of a nanolayer of ecDNA by plants on the surface of their leaves, stems, and roots could be a vital adaptation developed during evolution to increase the general fitness of a particular species [1].

Naked ecDNA, most of which is released as a result of cell death, is ubiquitous in the environment. Its concentration can reach up to 2 μg/L in soil, and up to 88 μg/L in natural aquatic environments [2], where it plays multiple intracellular and extracellular roles. At the intracellular level, it has been shown to be involved in functions including, horizontal gene transfer [3], providing nutrients [4]; and acting as a buffer to recruit or titrate ions or antibiotics [5]. At the extracellular level ecDNA has been shown to be a component of the biofilms of several bacterial species, where it plays different roles, such as regulating the attachment and dispersal of specific cell types in the biofilm [6], contributing to biofilm formation and providing physical strength and resistance to biological stress [7,8]. More recently, ecDNA has been documented to be associated with damage-associated molecular pattern (DAMP) that can cause species-specific plant damaged self-recognition [9,10].

In 2008, our laboratory discovered the insecticidal potential of contact unmodified antisense DNA (CUAD), opening a novel dimension for DNA-based plant protection research. CUAD biotechnology (CUADb or ‘genetic zipper’ technology) has emerged as an efficient and environmentally friendly platform for targeting hemipterans, thrips and mites. This approach leverages pest rRNA as a molecular target, significantly enhancing the efficiency and selectivity. Ribosomal RNA constitutes approximately 80-85% of the total cellular RNA, whereas mRNA accounts for only 5% [11]. Targeting mature rRNA and/or pre-rRNA increases the signal-to-noise ratio (∼10^5^:1; rRNA vs. random mRNA), improving specificity while reducing the likelihood of off-target effects. CUADb-based oligonucleotide insecticides, designed with species-specific sequences, exhibit selective efficacy and resistance avoidance by targeting conserved rRNA regions. On average, pest mortality rates range between 80 and 90% within 3-14 days following a single treatment with a 100 ng/ μL solution, resulting in the application of 1 mg of DNA per m^2^ of plant leaves [12]. This study underscores the potential of CUADb as an innovative and sustainable pest control strategy and highlights its parallels with naturally occurring extracellular DNA with insecticidal properties (eci-DNA), further supporting the role of extracellular DNA in plant defense mechanisms.

Oligonucleotide insecticides function through a conserved DNA containment (DNAc) mechanism, which can be triggered by exogenously applied antisense DNAs in insects. The DNAc mechanism operates in two steps: (1) arrest of the target rRNA, followed by its hypercompensation via rDNA transcription and (2) target rRNA degradation. The interaction between target rRNA and an oligonucleotide insecticide in the presence of DNA-guided rRNase, such as DNA-RNA hybrid-guided RNase H1, mimics a zipper-like mechanism facilitated by a DNA-RNA duplex (‘genetic zipper’ technology) [13]. This ‘genetic zipper’ mechanism, induces metabolic shifts toward lipid-based energy synthesis, enhancing ribosomes biogenesis and ATP production. Ultimately, wide-spread downregulation of kinases, including mTOR, which is a master regulator of ribosome biogenesis via mTORC1, causes a ‘kinase disaster’ (ca. 80% of kinases are down-regulated) due to ATP insufficiency, while significant RNase H1 upregulation occurs during DNAc in comparison with water-treated control [13].

The primary objective of oligonucleotide insecticides in agriculture is to produce crops that are free of organic xenobiotics. Although these insecticides may not serve as a universal solution at the moment, they could still be highly effective in controlling several dozen economically important insect pest species, representing a major advancement in sustainable plant protection. Current estimates suggest that CUADb-based ‘genetic zipper’ technology has the potential to effectively manage up to 50% of the most serious and insecticide-resistant pests on the planet using a simple and adaptable algorithm.

Recently, our group documented a naturally occurring insecticidal mechanism relying on unmodified DNA fragments [1,14]. However, whether RNA could have a similar effect is not yet known. Notably, RNA interference has a limited success in insect pest control, largely due to the poor understanding of its role in natural pest regulation mechanisms [15]. In our previous studies [1,13], we investigated the effects of eci-DNA fragments from the evergreen shrub *Pittosporum tobira* on the pest *Coccus hesperidum* revealing a pronounced insecticidal effect. Under natural conditions, eci-DNA concentrations in *P. tobira* leaves were measured at 24.37 ± 0.97 ng per cm^2^ [1]. Unexpectedly, we later found that eci-DNA is not exclusively composed of double-stranded fragments but also contains single-stranded DNA in the outermost layer of the leaf cuticle and is substantially enriched with chloroplast DNA. This study aims to address this critical knowledge gap highlighting a natural parallel to CUADb-based oligonucleotide insecticides and ‘genetic zipper’ technology in general.

## 2. Results

### 2.1. Eci-DNA released from P. tobira consists of short DNA fragments

Before testing the insecticidal activity of the released DNA, we isolated and purified eci-DNA from the leaf surfaces of *P. tobira*. Next, we loaded the sample onto a 1.8% agarose gel to check the size distribution. Unexpectedly, the size of eci-DNA was short and homogenous, with the majority of fragments being under 100 bp of DNA ladder (Figure 1).

**Figure 1.**
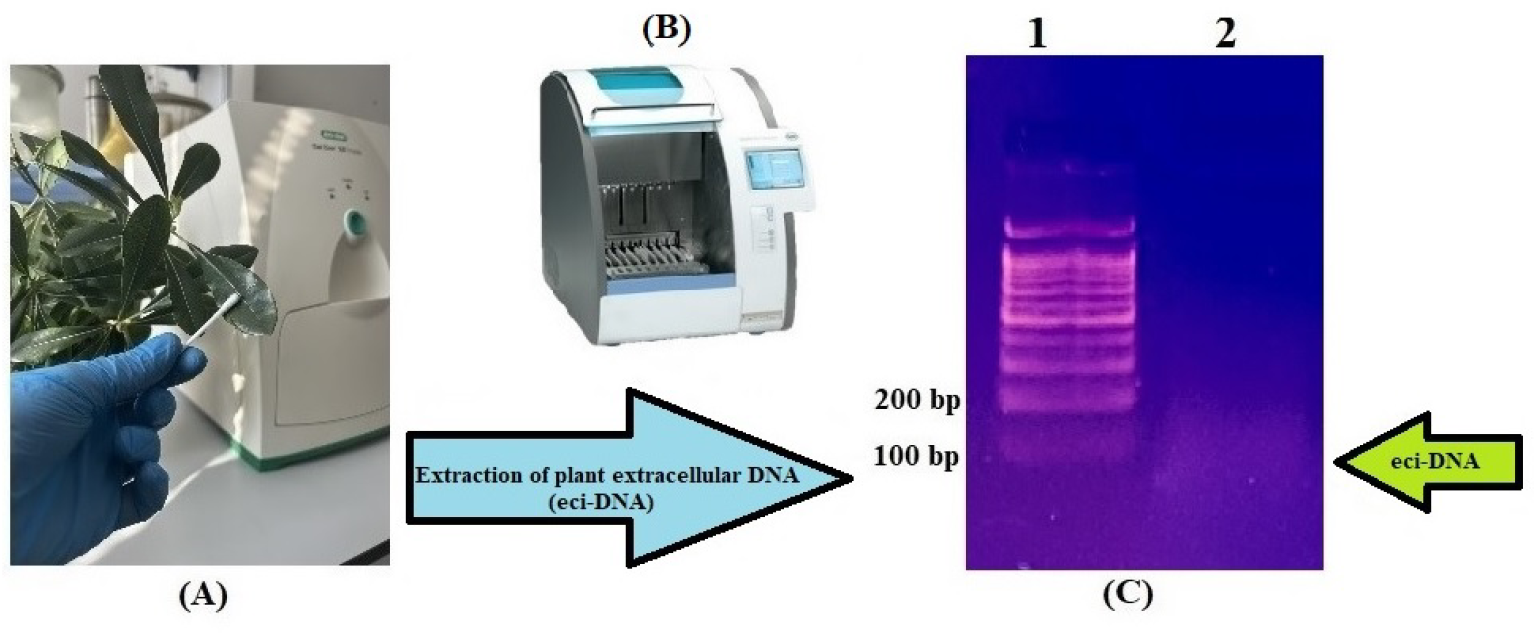
General scheme of collection and extraction of eci-DNA from the leaf surfaces of *P. tobira* (**A**) eci-DNA was collected from the leaf surfaces with wet cotton balls; (**B**) MagNA Pure Compact Instrument was used for the extraction of DNA; (**C**) Electropherogram (1.8 % agarose gel): 1 – DNA ladder 100 bp+; 2 – eci-DNA of *P. tobira* leaves.

This suggests that eci-DNA is a result of active fragmentation rather than being derived from dead and damaged cells, which would result in a DNA smear with different lengths, ranging from a few base pairs to mega-base pairs.

### 2.2. Eci-DNA shows pronounced insecticidal effect after topical application

Next, to check whether this uniformly secreted DNA could exhibit an insecticidal effect as an eci-DNA, we topically applied it to *C. hesperidum* larvae (1.2 ng of eci-DNA per larva of 2–2.2 mm^2^ body size). Four days post treatment, we assessed the mortality rate and found that eci-DNA killed approximately 80% of the larvae, while the scrambled control with random 56-mer single-stranded oligonucleotide (ACTG)_14_ did not show any significant toxicity when compared with the water-treated control group; both showed ca. 10% mortality (Figure 2).

**Figure 2.**
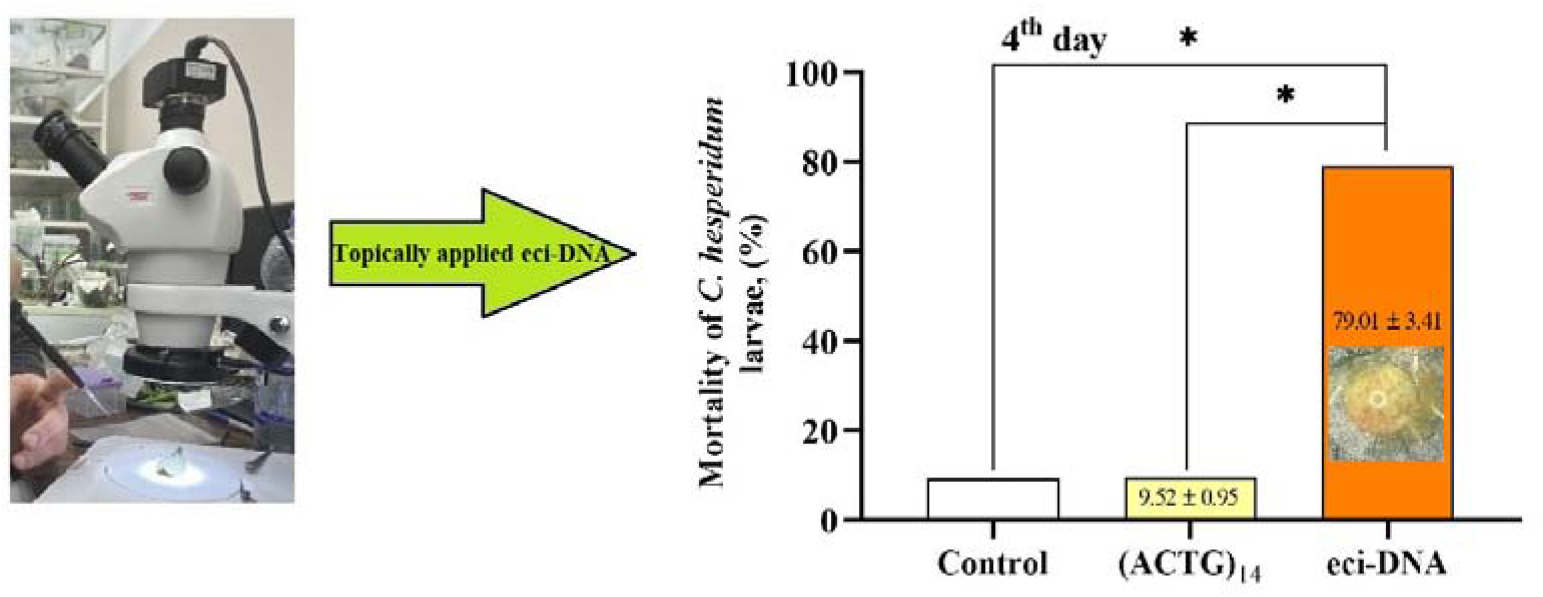
Insecticidal effect of eci-DNA on *C. hesperidum* larvae. Mortality of *C. hesperidum* larvae after topical treatment with eci-DNA, (ACTG)_14_ and water-treated control. The significance of the difference in the eci-DNA group compared to the (ACTG)_14_ and water-treated control is indicated by * at *p* < 0.01.

### 2.3. Comparison with synthetic technologies (‘genetic zipper’ technology)

The genomic DNA of insects is predominantly methylated [16,17]. Methyl groups make DNA less immunogenic to the organism itself [18] and also more resistant to DNA nucleases [19-21]. We studied the insecticidal activity of the methylated oligonucleotide insecticide Coccus(5mC)-11 (1 mg of DNA per m^2^ of leaves) and found its high insecticidal potential (76.82%) compared to unmethylated Coccus-11 (64.28%) on the 14^th^ day (Figure 3).

**Figure 3.**
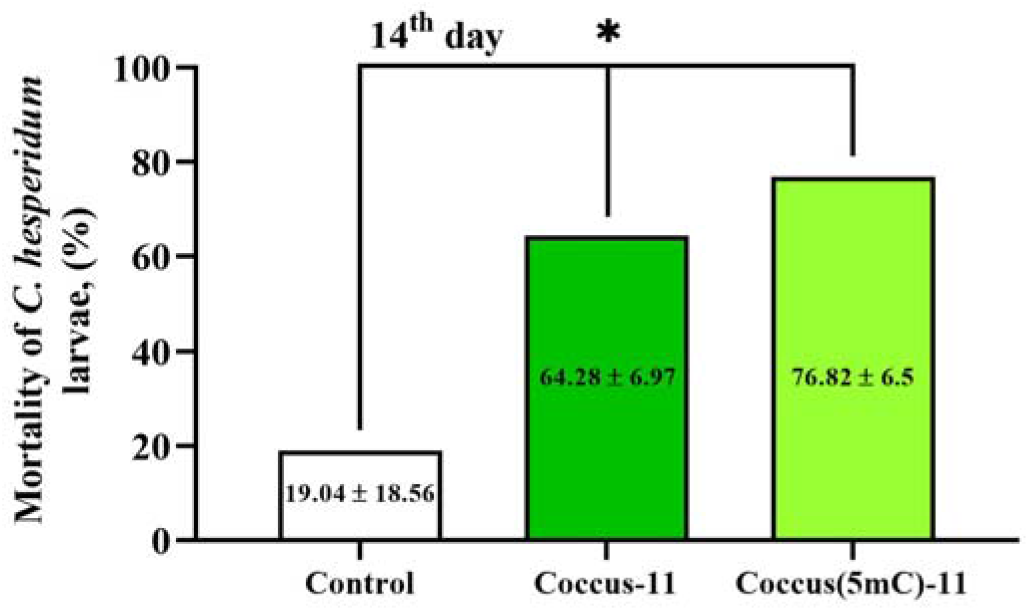
Insecticidal effect of unmethylated Coccus-11 (5′–CCATCTTTCGG–3’) and methylated Coccus(5mC)-11 (5′–(5mC)-(5mC)-AT-(5mC)-TTT-(5mC)-GG–3’) on *C. hesperidum* larvae. Mortality of *C. hesperidum* larvae after topical treatment with Coccus-11, Coccus(5mC)-11, and water-treated control. The significance of difference in the Coccus-11 and Coccus(5mC)-11 groups compared to the water-treated control is indicated by * at *p* < 0.01; experiments were carried out in the field, the average daytime temperature during the experiment was 16.27 ± 2.11 °C, at night – 13.07 ± 2.41 °C, and humidity was 76.94 ± 7.12%.

The oligonucleotide insecticide Coccus-11 has been well characterized in our earlier studies [1], has a high insecticidal potential and is a reliable reference oligonucleotide insecticide. The results indicate that DNA methylation increases its insecticidal potential (or at least does not reduce it) and methylated eci-DNA is also capable of possessing insecticidal effects on pests.

### 2.4. Eci-DNA mainly contains nuclear DNA

To understand the origin of eci-DNA and whether it is derived from nuclear or chloroplast DNA, we amplified both 5.8S rDNA (nuclear) and 23S rDNA (chloroplast) genes using specific primers. As a negative control, we measured the concentrations of nuclear rDNA and chloroplast rDNA in intact *P. tobira* leaves. Quantitative PCR results showed that the concentration ratio between nuclear rDNA and chloroplast rDNA was 7.7 to 1 in intact *P. tobira* leaves. Surprisingly, in the eci-DNA samples, the ratio between nucleus rDNA and chloroplast rDNA was 1.6 to 1, which represents approximately five-fold increase in chloroplast rDNA (Figure 4). This suggests that eci-DNA is substantially enriched by the chloroplasts DNA.

**Figure 4.**
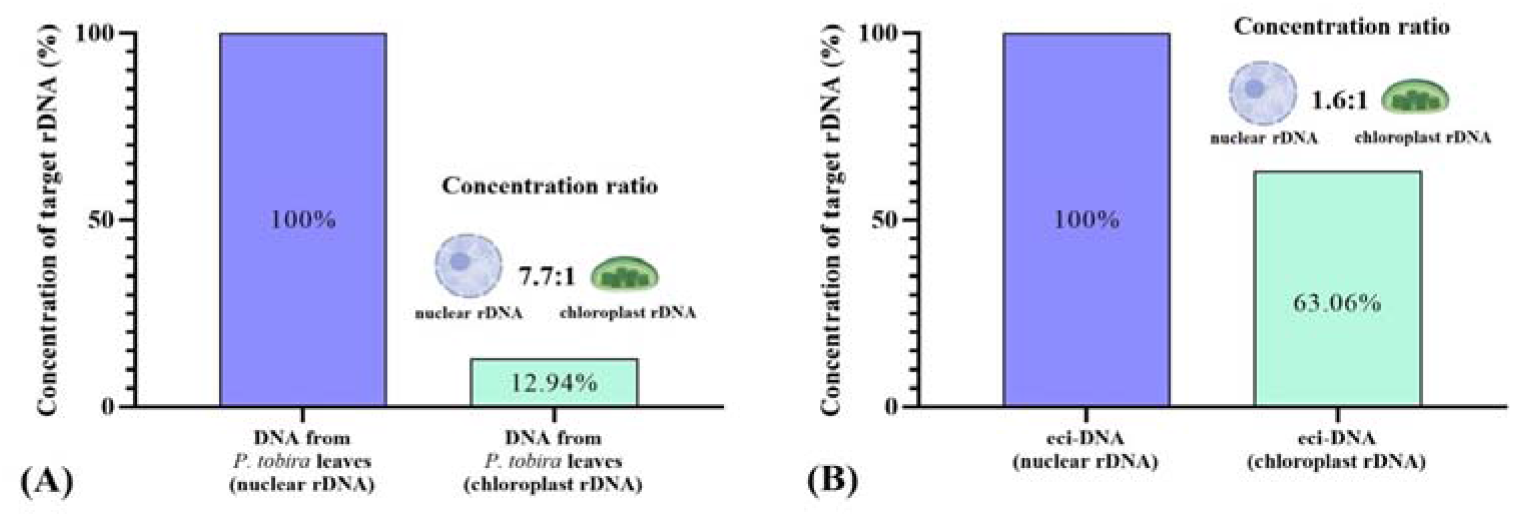
Relative concentration of nuclear rDNA and chloroplast rDNA in total DNA from intact *P. tobira* leaves (A) and eci-DNA (B). Nuclear rDNA was taken as 1 (100 %).

### 2.5. Eci-DNA is predominantly single-stranded

Next, we investigated the nature of the DNA fragments in eci-DNA, whether they were predominantly ssDNA or dsDNA. To this end, we first treated DNA extracted from intact *P. tobira* leaves with ExoI nuclease (degrades ssDNA), which specifically cleaves ssDNA, followed by qPCR quantification. We found that approximately 99% of the DNA was double-stranded in both the nuclear and chloroplast fractions (Figure 5).

**Figure 5.**
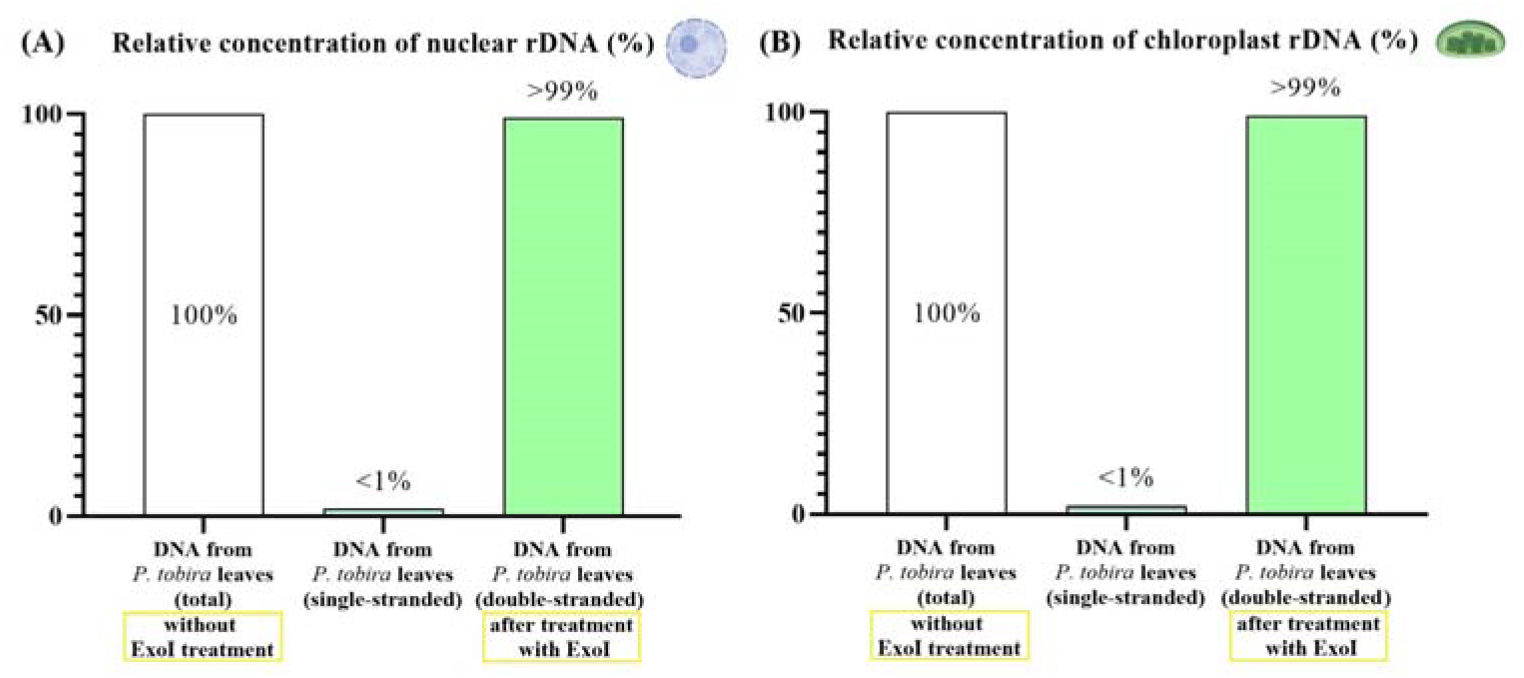
Relative concentrations of nuclear rDNA (A) and chloroplast rDNA (B) in intact *P. tobira* leaves after ExoI nuclease treatment. Total DNA from intact plant leaves (non-treated with ExoI nuclease) was set to 1 (100 %).

Surprisingly, eci-DNA treatment with ExoI nuclease completely reversed the previous profile and showed that it mainly composed of single-stranded DNA: 24.8:1 for nuclear DNA and 103.1:1 for chloroplast DNA (Figure 6).

**Figure 6.**
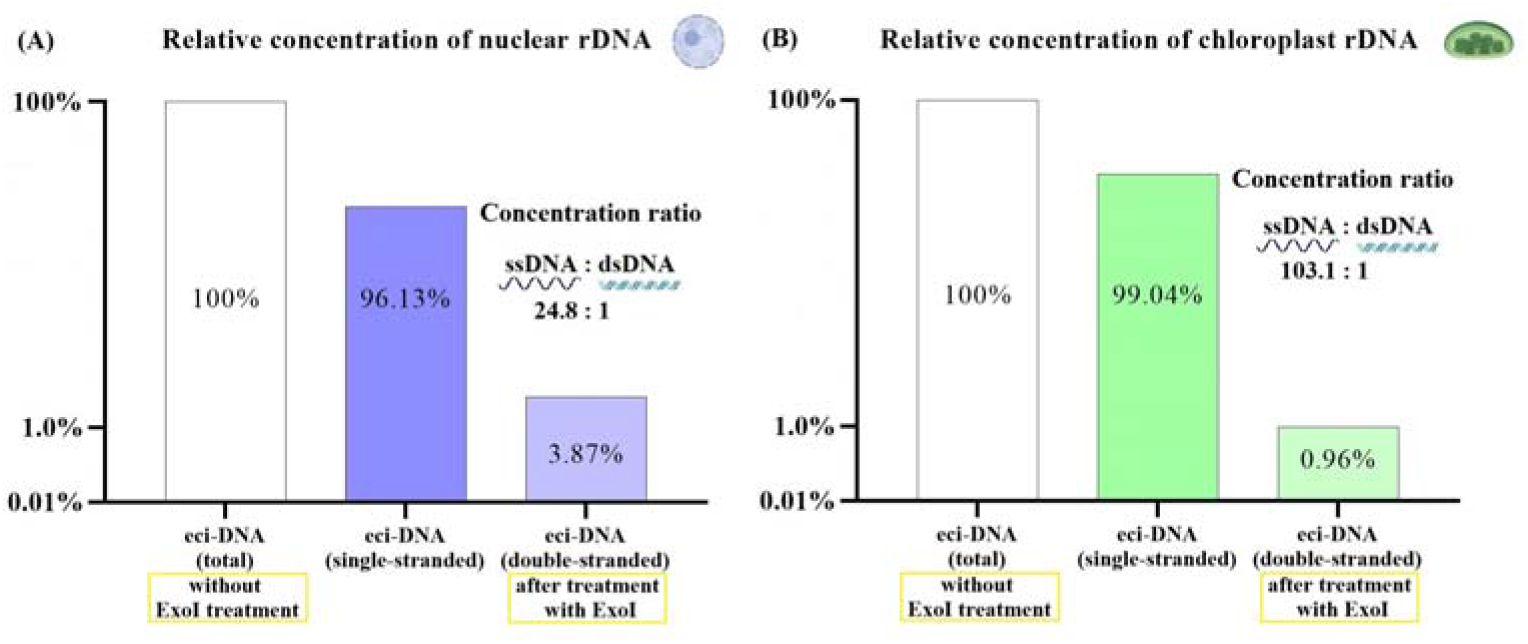
Relative concentrations of nuclear rDNA (A) and chloroplast rDNA (B) from eci-DNA after treatment with ExoI nuclease. Eci-DNA (non-treated with ExoI nuclease) was set to 1 (100 %).

### 2.6. Pittosporum tobira leaves show a positive staining for eci-DNA

To confirm the presence of eci-DNA in *P. tobira* leaves, we also performed histological analyses in which we stained both ssDNA and dsDNA in different parts of the leaves. In the control section (without staining), several structures of the anatomical preparation of the leaf exhibited autofluorescence (Figure 7, A).

**Figure 7.**
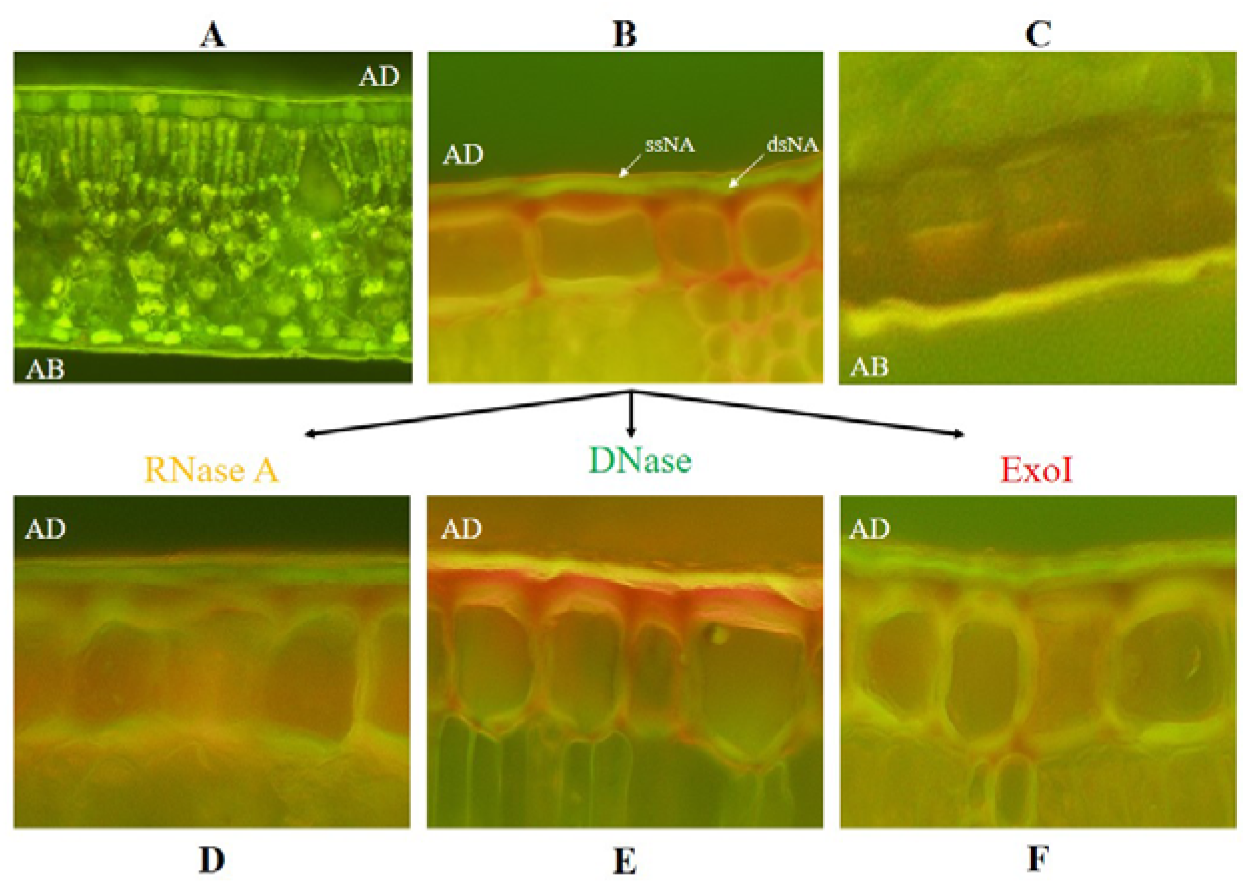
Anatomical preparations of *P. tobira* leaf cross-sections stained with acridine orange fluorochrome: A – temporary preparation in native state, non-treated; B – preparation stained with fluorochrome (adaxial surface, AD): ssNA (red color) – layer of single-stranded nucleic acid, dsNA (green color) – layer of double-stranded nucleic acid (explanation is in the text).; C – preparation stained with fluorochrome (abaxial surface, AB); D – treated with RNase A before staining with fluorochrome; E – treated with DNase before staining with fluorochrome; F – treated with ExoI before staining with fluorochrome

However, after staining with fluorochrome acridine orange, we found that the cuticle on the adaxial part of the leaf fluoresced predominantly in green, indicating the presence of double-stranded nucleic acid. A thin layer of the peripheral part of the cuticle fluoresced clearly in red-orange, confirming the presence of single-stranded nucleic acid in this layer (Figure 7, B). Interestingly, the *C. hesperidum* pest predominantly settled on the abaxial part of *P. tobira* leaves, where the nucleic acid layer was less pronounced (Figure 7, C) and was mainly yellow. Further studies were conducted to determine which nucleic acid (DNA or RNA) was present and fluoresced on the leaf surface in the layers fluorescing green and red-orange. For this purpose, before staining with fluorochrome, *P. tobira* leaves were pre-treated for 20 min with RNase A (Figure 7, D), DNase (Figure 7, E), and ExoI (Figure 7, F) nucleases and then washed out with water.

The results revealed that treatment of the adaxial side of leaves with RNase A (which degrades ssRNA and dsRNA) did not induce noticeable differences from the control group, indicating the absence of both ssRNA and dsRNA on the surface of *P. tobira* leaves. The green and red-orange cuticle layers remained unchanged (Figure 7, D). In contrast, under DNase treatment (degrades ssDNA and dsDNA), the green fluorescence of the cuticle on the adaxial side of the leaf changed to yellow (similar in color to abaxial surface (AB) of leaves where the visible nucleic acid layer on cuticle was not detectedb), confirming that the double-stranded nucleic acid found on the leaves was dsDNA. The clearly structured red-orange layer of the single-stranded nucleic acid also disappeared, indicating that it was formed by ssDNA. These effects led to the formation of a loose layer characteristic of unstructured and partially degraded ssDNA (Figure 7, E). Upon ExoI treatment (which degrades ssDNA), the red-orange layer of the single-stranded acid became noticeably thinner, confirming the presence of single-stranded DNA in the outermost layer of the leaf surface (Figure 7, F). As expected, the green layer of double-stranded nucleic acid remained unchanged under action of ExoI. The results obtained were consistent with those obtained by real-time PCR and confirmed the presence of single-stranded DNA in the outermost layer of the cuticle of *P. tobira* leaves. It should be noted that ssNA is present in great quantities in epidermal layer of cells (Fig. 7, B; red color around epidermal cells). Clearly, epidermal cells actively participate in the formation of ssNA and ssDNA, in particular, which then penetrate the cell surface.

### 2.7. Differential gene expression (DGE) analysis of C. hesperidum reveals activation of retrotransposons caused by Coccus-11 oligonucleotide insecticide

The data obtained from the DGE analysis indicated a significant activation of retrotransposons in *C. hesperidum* cells under the influence of the oligonucleotide insecticide Coccus-11. The expression of the reverse transcriptase domains (RVT1 and RVT2), ribonuclease H domain, along with integrase and peptidase, of retrotransposons (mainly Pao and Tf2), significantly increased (Figure 8). Pao retrotransposons potentially act as molecular machinery for antiviral defense [22,23], whereas Tf2 retrotransposons impact the rapid adaptive stress responses of insects (such as insecticide application or host-plant resistance) [24].

**Figure 8.**
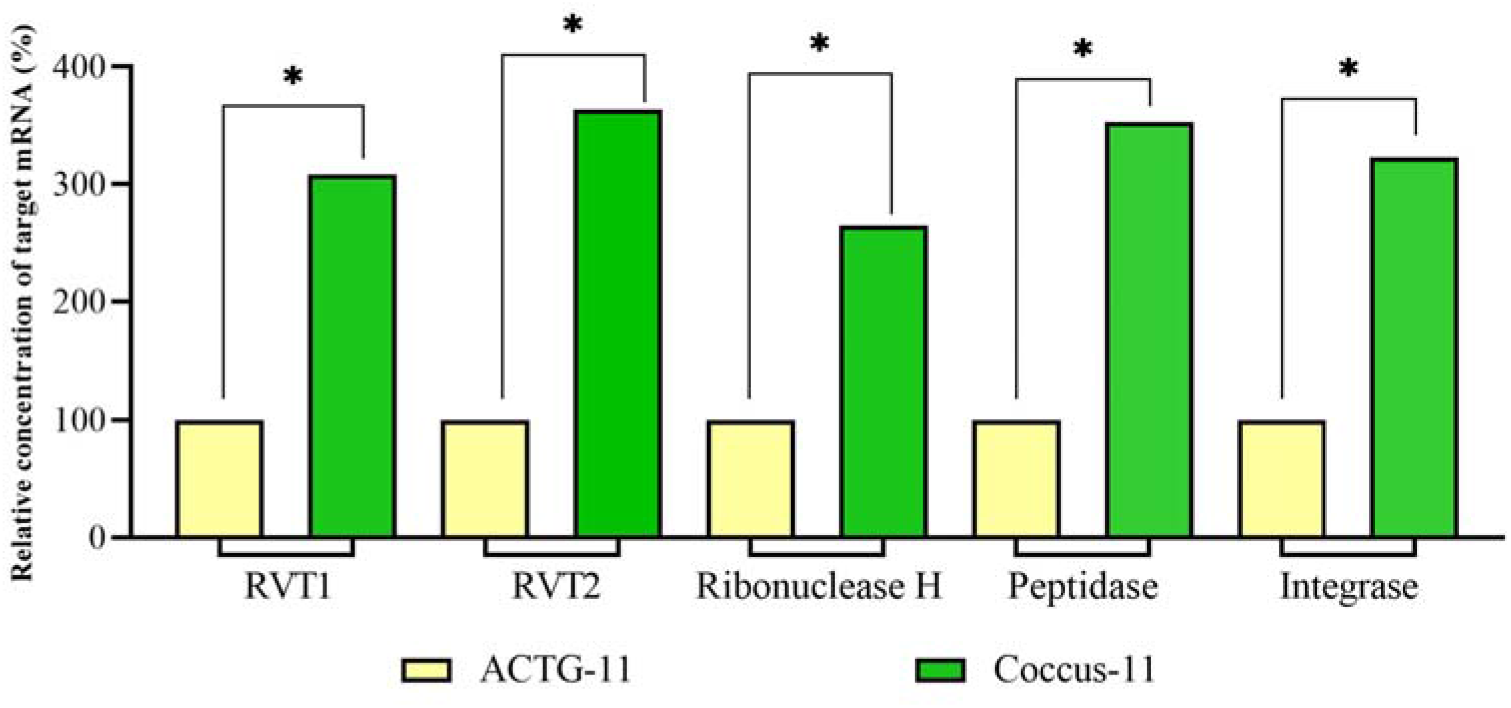
Relative expression of the reverse transcriptase domains (RVT1 and RVT2), ribonuclease H domain, peptidase, and integrase of retrotransposons. The significance of difference in the Coccus-11 group compared to the ACTG-11 is indicated by * at *p* < 0.05; expression in ACTG-11 group was set to 100%

Accordingly, the expression of key inhibitors of retrotransposon replication: HSP90AA1, HSP90BB1, Aubergine, and AGO2 significantly decreased (Figure 9).

**Figure 9.**
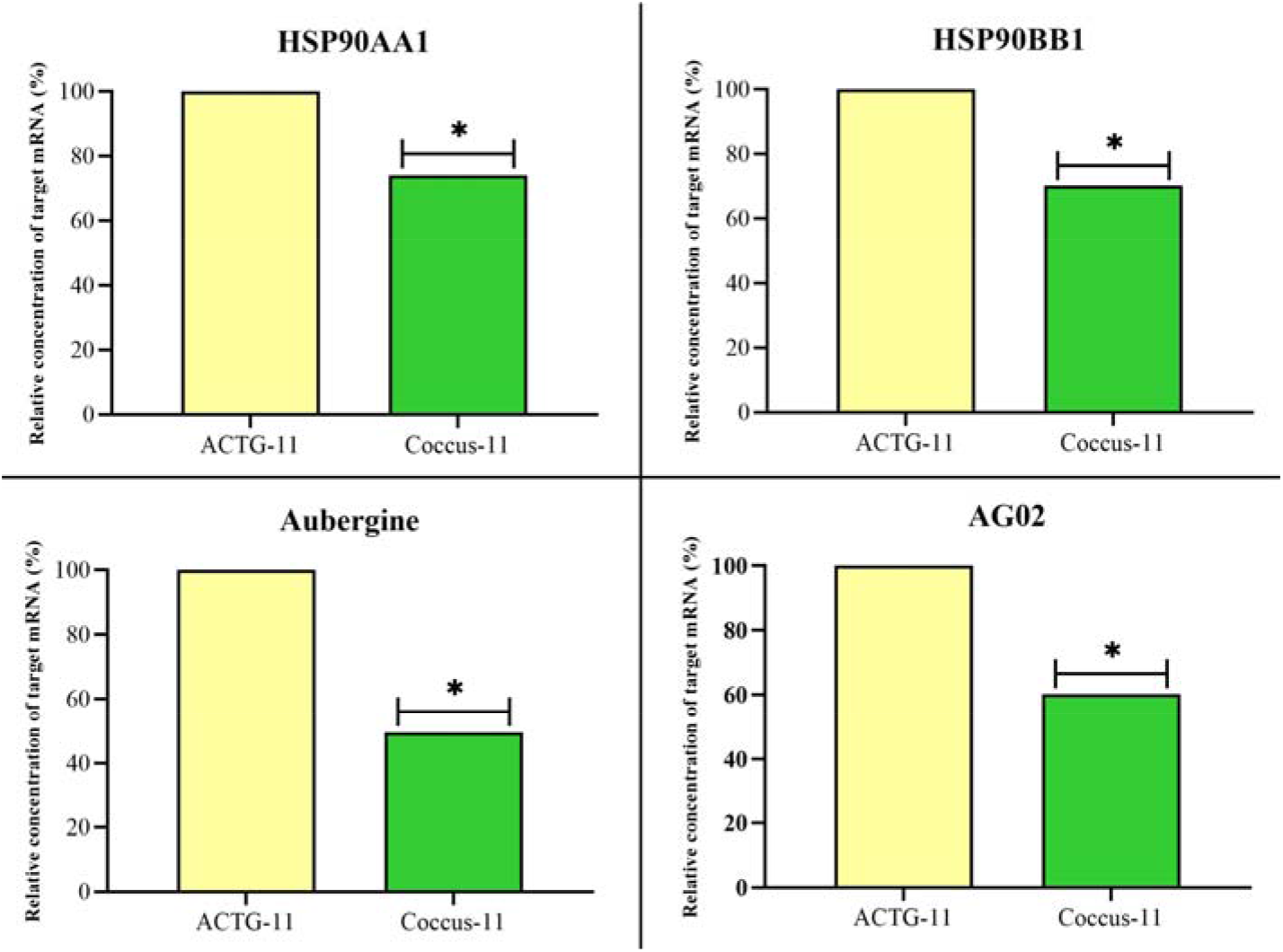
Relative expression of key inhibitors of retrotransposon replication: HSP90AA1, HSP90BB1, Aubergine, and AGO2. The significance of difference in the Coccus-11 group compared to the ACTG-11 is indicated by * at *p* < 0.05; expression in ACTG-11 group is taken as 100%

These inhibiting proteins act as part of the host defense system, specifically within the PIWI/piRNA pathway [25-27] while the latter, AGO2, plays a critical role in inhibiting retrotransposon replication, primarily collaborating with the PIWI/piRNA pathway to silence retrotransposons [28-30]. While PIWI proteins are central to germline expression, AGO2 is often involved in both somatic and germline tissues by utilizing endogenous siRNAs to target and cleave retrotransposon transcripts [31-33].

## 3. Discussion

The results obtained in this study suggest that *P. tobira* cells possess a mechanism for fragmenting genomic DNA into smaller fragments and converting them into single-stranded DNA (ssDNA). Similar to purpose-driven ‘genetic zipper’ technology-based oligonucleotide insecticides, eci-DNA likely functions via evolution-driven targeted gene silencing (Table 1).

**Table 1.** Comparative overview of ‘genetic zipper’ technology and extracellular DNA with insecticidal properties (eci-DNA).

| Key Features | eci-DNA | ‘genetic zipper’ technology |
| --- | --- | --- |
| Effector | dsDNA << ssDNA | ssDNA |
| Length | 50-100 nt | 11-20 nt |
| Target site | Whole pool of cell RNA | Pre-rRNA and rRNA |
| Mechanism of action | DNA containment | DNA containment |
| Production | Leaves | Synthetic |
| Target pest | Broad spectrum | Species-specific |
| Development | Evolution-driven | Purpose-driven |

Observed phenomenon is particularly intriguing in the context of eci-DNA, which appears to modulate gene expression in *C. hesperidum* larvae through sequence complementarity to their mRNAs and non-coding RNAs, including rRNAs, causing pest death. This is supported by the discovery of two 11 nucleotides (nt) sequences in the plastome of *P. tobira* that exhibit perfect complementarity to *C. hesperidum* rRNA (28S, MF594311.1; 18S, MF594274.1), suggesting a potential mechanism for DNA containment-mediated insecticidal effects [13]. Given that the rDNA of *C. hesperidum* is only partially sequenced, additional complementary sequences may exist, further supporting this hypothesis. Thus, eci-DNA can also act by targeting pest rRNAs in a manner similar to that of CUADb-based oligonucleotide pesticides [12]. It should be noted that eci-DNA in agarose gel was purified from impurities twice, using specialized nucleic acid purification kits: 1. ColGen kit (Syntol, Moscow, Russia) — was used for denaturation of proteins in DNA-protein complexes using a chaotrope followed by adsorption of DNA on a silica column membrane, and 2. Isolation DNA kit (Roche Diagnostics GmbH, Mannheim, Germany) — denaturation of proteins in DNA-protein complexes using a chaotrope followed by DNA purification on magnetic particles. It is most unlikely that any other substance (such as proteins) besides nucleic acids could have an insecticidal effect after eci-DNA purification.

In our opinion, plant single-stranded methylated (5mC) eci-DNA can also serve as a complex of primers for chain extension on complementary mRNA and rRNA templates within insect cells. For this reason, the most critical role will be played by the 3’-end of the DNA sequence, which should be complementary to the RNA target, based on literature data, by at least 6 nt to trigger chain elongation [34]. Moreover, the results indicate that DNA methylation increases its insecticidal potential (or at least does not reduce it) and methylated eci-DNA is also capable of possessing insecticidal effects on pests. We assume that, during the DNA containment mechanism, DNA elongation (signal amplification) based on eci-DNA can be complemented in some insects, e.g. *C. hesperidum*, by RT-RNase H of retrotransposons [35]. Such response of the cells could be explained by their pursuit to strengthen (amplify) the signal and obtain a longer DNA strand that can find more parts of the target RNAs for their degradation by RNase H (obviously existing also as a defense against viruses). This is probably the reason behind the importance of the complementarity of the oligonucleotide insecticides’ 3′-end in the insecticidal effect in scale insects, because 3′-end is important for further DNA synthesis in the 5′-3′ direction [36]. To further investigate this idea, we found that under the influence of antisense oligonucleotide Coccus-11, retrotransposons (Pao and Tf2) were activated with the formation of a significant number of reverse transcriptase (RT)-ribonuclease H (RNase H) enzyme units. RT and RNase H are among the most ancient and most abundant protein folds [37] and are crucial for retrotransposition, often acting as a multifunctional enzyme where RNase H degrades RNA within DNA/RNA hybrids during DNA synthesis by RT [37-39]. Retrotransposons constitute a massive portion of insect genomes acting as major drivers of genome evolution and adaptation; and shaping genome size and structure [40]. Retrotransposition regulates gene expression, aids stress responses, and facilitates pathogen resistance. Although often suppressed by the host PIWI/piRNA pathway to prevent excessive damage, these elements are frequently ‘rehabilitated’ by the genome to serve different functional roles [41,42]. Moreover, retrotransposons are known to insert into the ribosomal DNA (rDNA) of insects targeting a highly conserved site within the 28S rRNA genes [43]. While commonly viewed as genomic parasites that disrupt rRNA function, recent research indicates that in the germline retrotransposons can act as a ‘mutualistic’ element by creating double-stranded DNA breaks that stimulate the expansion of rDNA copy numbers, crucial for maintaining fertility across generations [44].

The obtained results indicate that retrotransposon RT-RNase H is an ‘accomplice’ of oligonucleotide insecticides in the cell, which not only cleaves the target 28S rRNA [13], but it also capable of increasing the length of the oligonucleotide insecticide Coccus-11 (and strengthens insecticidal signal) with the help of RT domain on the rRNA template and its subsequent degradation by the RNase H domain. Thus, not only RNase H1 [13], but also RT-RNase H of retrotransposons is capable of contributing to the degradation of target rRNA under the action of oligonucleotide insecticides, such as Coccus-11. In the second step of the DNA containment mechanism [13,36] (target RNA degradation), the key enzymes are DNA-guided rRNases, such as DNA-RNA hybrid-guided RNase H1 [13]. This study shows that DNA-guided RT-RNase H of retrotransposons can significantly contribute to the degradation of the target rRNA under the influence of an oligonucleotide insecticides. Moreover, RT-RNase H was the key enzyme that is significantly upregulated in *C. hesperidum* cells compared to the random primer ACTG-11 (5′–ACT–GAC–TGA–CT–3′). On the contrary, RNase H1 was not significantly up-regulated in Coccus-11 group in comparison with random primer ACTG-11 (p>0.05). Therefore, it is apparently due to the impact of RT-RNase H, not RNase H1, that we observe in most cases a more rapid decrease in the concentration of target rRNA under the influence of oligonucleotide insecticides compared to random oligonucleotides [13] and oligonucleotide insecticides with single nucleotide substitutions [36,45]. Most of our studies using oligonucleotide insecticides were conducted on representatives of the order Hemiptera, whose genomes have, on average, a higher absolute retrotransposon content per cell than those of other insect orders [40,46]. In other words, oligonucleotide insecticides (DNA insecticides) and eci-DNA can be considered as ‘primers’ for large-scale reverse transcription in cells of hemipterans and the subsequent degradation of the target RNA by ubiquitous RT-RNase H (Figure 10).

**Figure 10.**
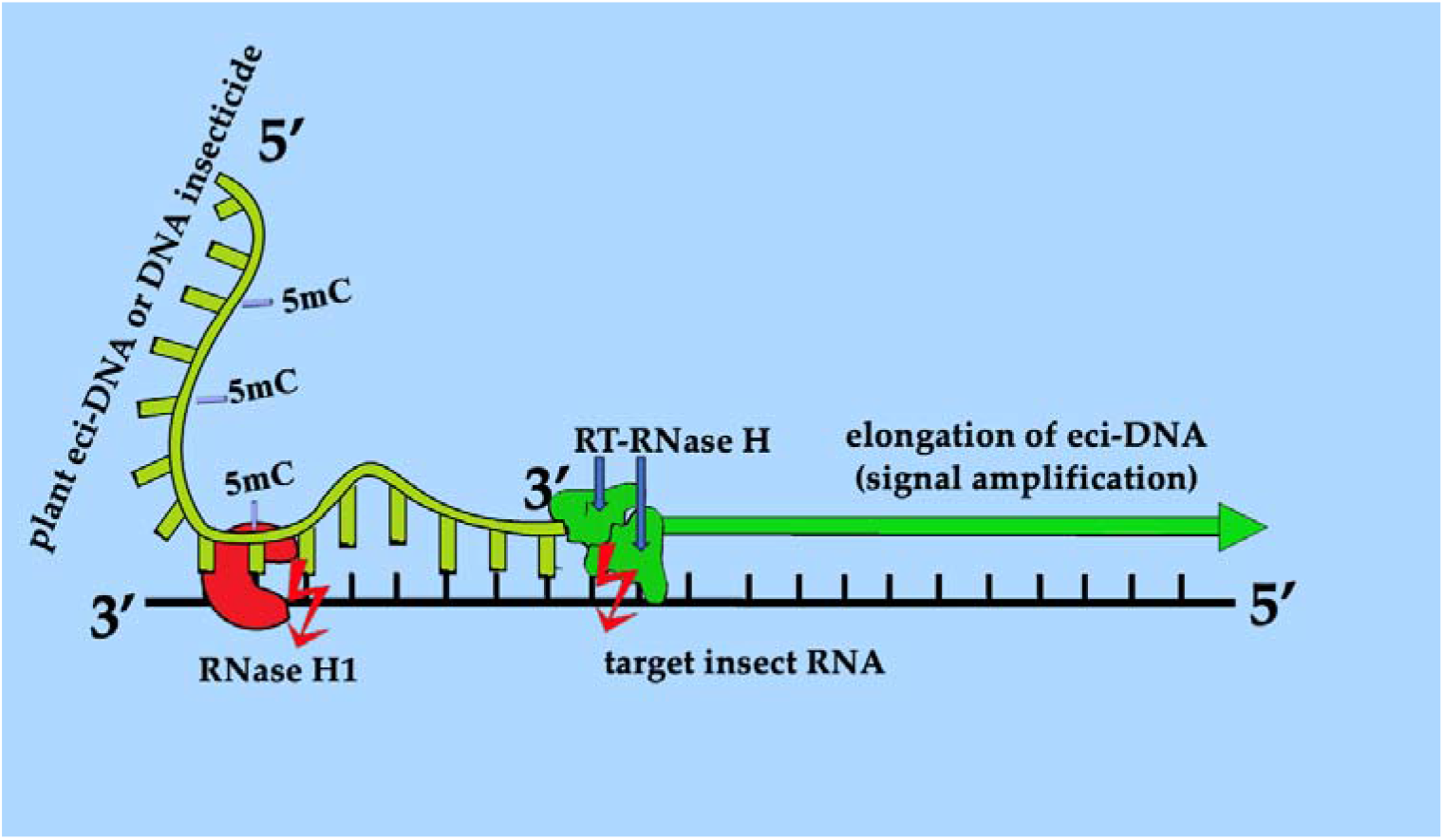
Proposed mechanism of target RNA degradation by eci-DNA or DNA insecticide in insect cells: RT-RNase H elongates plant eci-DNA or DNA insecticide (signal amplification) and enhances the degradation of target insect RNA by the RNase H module of RT-RNase H and RNase H1.

In this study, we established for the first time a link between the use of the unmodified antisense DNA oligonucleotide Coccus-11 and the activation of retrotransposons (deemed as ‘retrotransposon carnival’), which has both practical and fundamental significance. From a practical standpoint, this discovery will help improve oligonucleotide insecticides and ‘genetic zipper’ technology in general, expanding the capabilities of this approach to those insect groups whose genomes contain higher copies of retrotransposons per insect cell. From a fundamental standpoint, this finding brings us closer to understanding the phenomenon of horizontal gene transfer, for example, between insects and plants [47]. Extracellular DNA from various organisms is present in significant quantities in nature, which can be absorbed by them [48] and cause various effects [49-52], including insecticidal effects [1]. The ‘retrotransposon carnival’ demonstrated in this study can facilitate horizontal gene transfer using antisense rDNA in germline cells. The activity of RT-RNase H culminates in the formation of dsDNA, which is capable of homologous recombination [37,53]. Thus, if a single-stranded DNA fragment of any plant gene from eci-DNA enters an insect cell and initiates reverse transcription via RT-RNase H on any insect RNA using its 3’-end, the resulting dsDNA can integrate into the region of the insect genome with the highest homology. Subsequently, for any plant gene, due to the overlap of single-stranded DNA fragments in eci-DNA (because in each cell, cuts in the plant genomic DNA will occur primarily in different sites), the next newly formed dsDNAs will integrate into the same region of the insect genome where fragments of the given plant gene are already embedded. Thus, step by step, a new plant gene is formed within the insect genome, which, over time, after refinement by intrachromosomal rearrangements and point mutations, can become a functionally active plant gene within the insect genome. Organisms whose extracellular DNA frequently contacts each other (for example, a plant and its insect pest) have a greater chance of horizontal gene transfer than a random pair of organisms in any given ecosystem.

The obtained results indicate that DNA containment (DNAc) [13,36] is a unique mechanism of action of oligonucleotide insecticides (Table 1). In the first step, the target rRNA is functionally ‘arrested,’ leading to its hypercompensation. In the second step, the target rRNA undergoes degradation mediated by DNA-guided rRNases such as RT-RNase H and RNase H1 [13]. Generally, all oligonucleotide insecticides and random DNA oligonucleotides initiate hypercompensation of rRNA [13], but subsequent substantial rRNA degradation and insect mortality occur only when an oligonucleotide perfectly matches the target rRNA [12,13,36,54]. Obviously, RT-RNase H plays a key role in DNAc and the insecticidal effect of antisense oligonucleotides, along with ATP depletion and ‘kinase disaster’. Many experimental data suggest that imperfectly complementary (random) oligonucleotides do not efficiently recruit DNA-guided rRNases, such as RT-RNase H and RNase H1, and insect cells ultimately restore homeostasis by degrading random oligonucleotides [13].

A striking observation in our study was the shift in the ratio of nuclear to chloroplast rDNA in eci-DNA compared to intact leaf DNA. While intact *P. tobira* leaves exhibited a nuclear-to-chloroplast rDNA ratio of 7.7:1, this ratio shifted to 1.6:1 in eci-DNA, indicating a substantial enrichment of chloroplast-derived DNA. This finding suggests that *P. tobira* may selectively incorporate chloroplast DNA into eci-DNA. One plausible explanation is that chloroplasts, which are essential for photosynthesis, may become less vital in cells covered by sedentary insect pests due to the accumulation of honeydew [55,56]. Deprived of direct sunlight, these cells may repurpose chloroplast DNA to produce eci-DNA as a defensive adaptation including chloroplast DNA. This would be a novel adaptation, allowing plants to utilize existing cellular components for pest defense without significant metabolic trade-offs. This hypothesis aligns with previous reports on chloroplast degradation in senescing or stressed plant cells [57], suggesting a broader role for plastid dynamics in plant-insect interactions. Additionally, our results indicate that outermost layer of eci-DNA predominantly consists of ssDNA, as confirmed by nuclease treatment and real-time PCR analysis. The conversion of plant-derived DNA into ssDNA with insecticidal properties (eci-DNA) may enhance its stability or facilitate its interaction with pest nucleic acids, thus increasing its effectiveness as an insecticidal agent [58]. It remains unclear whether this ssDNA formation is an active regulatory process by the plant or a consequence of enzymatic degradation by microbial nucleases. Further research is needed to determine whether different environmental or biotic factors influence this ssDNA predominance.

The ecological implications of eci-DNA extend beyond *P. tobira* and *C. hesperidum*. Given the widespread occurrence of sternorrhynchans (including scale insects, mealybugs, and psyllids) in tropical and subtropical regions, it is conceivable that eci-DNA-mediated pest control is a common strategy in evergreen plants. This study provides new insights into a naturally occurring defense mechanism that parallels synthetic ‘genetic zipper’ technology. Of note, similar to our findings on insecticidal role of ssDNA, particularly plastidic ssDNA in eci-DNA, represented by our team in preprint in 2025 (https://doi.org/10.1101/2025.11.18.689116, accessed on 26 February 2026) and in this article, were found by researchers studying dsRNA. They identified a plastidic dsRNA (dsNode343) in oilseed rape which contributes to defense against the cabbage stem flea beetle and published their results in preprint in 2026 [59]. Future investigations should explore whether eci-DNA production varies among plant species, whether different pests elicit distinct DNA fragment compositions, and whether environmental conditions influence eci-DNA efficacy. Overall, our findings suggest that eci-DNA represents a previously unrecognized form of plant defense, where selective DNA fragmentation and chloroplast DNA enrichment play a role in mitigating herbivory by sedentary insect pests. Understanding the molecular underpinnings of this process could pave the way for novel bioinsecticidal applications, leveraging natural plant defense mechanisms for sustainable pest management similar to ‘genetic zipper’ technology.

It is important to note that this study examined the outermost layer of eci-DNA of the plant itself on the leaf cuticle. We found that plant eci-DNA is single-stranded and enriched in chloroplast DNA compared to genomic DNA in intact *P. tobira* cells. However, we understand that microbial and insect DNA may also contribute to the formation of eci-DNA. The form in which their DNA may present in eci-DNA is still unknown and will be the subject of further research.

## 4. Materials and Methods

### 4.1. Origin of P. tobira and C. hesperidum

We identified *P. tobira* Thunb. (Apiales: Pittosporaceae) with *C. hesperidum* L. larvae (Hemiptera: Coccidae) on its leaves at the Nikita Botanical Garden (Yalta, Republic of Crimea, Russia).

### 4.2. Collection of eci-DNA from P. tobira leaves

Eci-DNA was collected during summer from the adaxial (AD) and abaxial (AB) surfaces of around 2400 *P. tobira* leaves gently using wet cotton swabs and filtered through a 0.2-micron syringe filter (Whatman, Maidstone, UK) (Figure 1). The total 10 mL filtrate was concentrated to 400 μL under vacuum (Heidolph, Schwabach, Germany) and used for subsequent experiments.

### 4.3. Gel electrophoresis

To assess the quality and consistency of extracted eci-DNA, 25 μl of filtrate was loaded per well onto a 1.8 % agarose gel, stained with 1% ethidium bromide, and subjected to electrophoresis in Tris-borate-EDTA (TBE) buffer at 10 V/cm for 45 min. The primary eci-DNA fraction was approximately 100 bp in length, as determined using a 100 bp+ DNA ladder (marker) (Figure 1).

### 4.4. Purification of eci-DNA from agarose gel

Eci-DNA was extracted from agarose gel using the ColGen kit (Syntol, Moscow, Russia). Gel fragments containing DNA were excised using a sterile scalpel and transferred to 1.5 mL Eppendorf tubes. The gel mass was determined, and a 3:1 volume of binding buffer was added (e.g., 600 μL for 200 mg of gel). The mixture was incubated at 55 °C for 10 minutes, until complete dissolution. Purification was performed using the Isolation DNA kit (Roche Diagnostics GmbH, Mannheim, Germany) and MagNA Pure Compact Instrument (Roche Diagnostics GmbH, Mannheim, Germany). The purified DNA was concentrated to 30 μL under vacuum (Heidolph, Schwabach, Germany) (Figure 1).

### 4.5. Nuclease Treatment

The RNase A (Vazyme Biotech, Nanjing, China), DNase (Roche, Mannheim, Germany), and exonuclease I (ExoI) (GenTerra, Moscow, Russia) were used according to the manufacturer’s instructions. For DNase treatemnt, 10 μL of eci-DNA was mixed with 3 μL of DNase in the buffer. For ExoI treatment, 10 μL of eci-DNA was mixed with 1 μL of ExoI and 2 μL of 1X ExoI buffer. ExoI nuclease treatment was used to calculate percentage of single-stranded DNA in eci-DNA during real-time PCR. Both reactions were incubated at 37°C for 15 minutes followed by enzyme inactivation at 80°C for 15 minutes. RNase A, DNase and ExoI were topically applied to *P. tobira* leaves by placing a 20 μL droplet on the adaxial side of the leaf and gently distributed by pipette tip.

### 4.6. PCR analysis of P. tobira eci-DNA

*Pittosporum tobira* eci-DNA was analysed using real-time PCR with the fluorescent dye SYBR Green I. The reaction mixture contained 3 μL of DNA, 5X qPCRmix-HS SYBR master mix (Evrogen, Moscow, Russia) and specific primers (Table 2). PCR amplification was performed on a LightCycler^®^ 96 instrument (Roche, Basel, Switzerland) using the following thermal cycling conditions: an initial denaturation at 95°C for 10 minutes, followed by 45 cycles of denaturation at 95°C for 10 seconds, annealing at 54°C for 15 seconds, and elongation at 72°C for 10 seconds. Each reaction was performed in triplicate. Melt curve analysis was conducted to confirm amplification specificity and detect potential non-specific products.

**Table 2.**
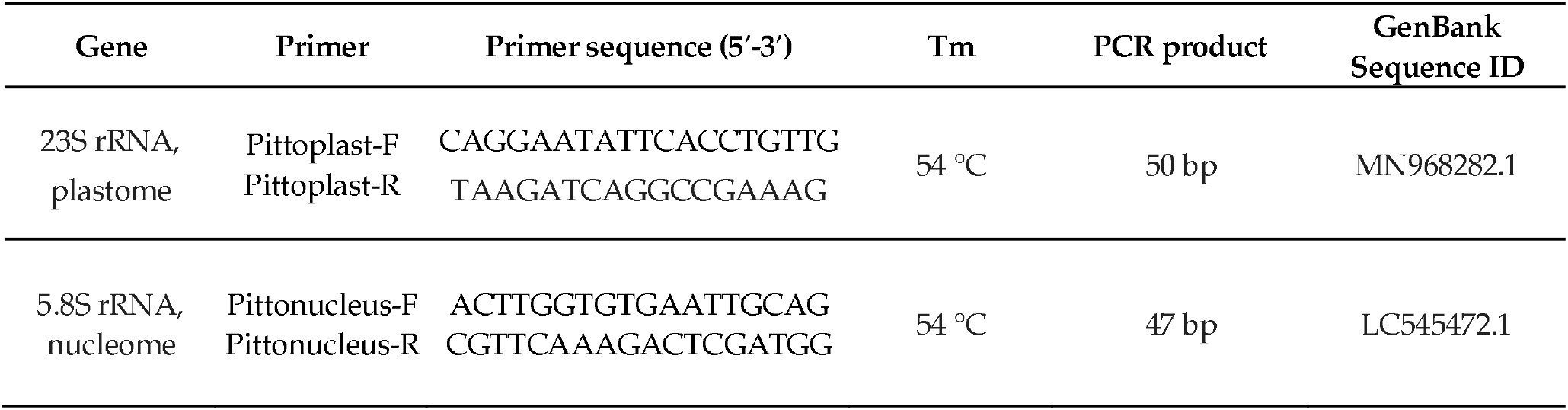
Primer sequences for real-time PCR using SYBR Green I system.

### 4.7. Application of eci-DNA as a contact insecticide in lab contidions

Eci-DNA (1.2 ng/μL in nuclease-free water) was topically applied to *C. hesperidum* larvae by placing a 1 μL droplet on the dorsal surface of each larva (average body size: 2–2.2 mm^2^) using a pipette. As a scrambled control of ssDNA fragment, we used 56 base nucleotide long random primer (containing 14 consecutive ACTG sequences (ACTG)_14_) in the same concentration. The control group received nuclease-free water. Experiments were conducted in triplicate, with ca. 100 larvae per treatment group, which were included in the statistical sample to assess the survival rate. Mortality was calculated as the percentage of dead larvae relative to the total number of individuals treated. Insects were counted using a Nikon SMZ745 microscope (Nikon Inst., Gyoda, Japan) and images were captured with a Toupcam-UCMOS-05100KPA camera (Asmetec, Kirchheimbolanden, Germany).

### 4.8. Application of oligonucleotide insecticides (Coccus-11 and Coccus(5mC)-11) in field conditions

Oligonucleotide insecticide sequences were designed based on the 28S rRNA sequence of *Coccus hesperidum* (isolate S6A395) retrieved from the GenBank database (https://www.ncbi.nlm.nih.gov/nuccore/MT317022.1, accessed on 30 March 2025). Two short oligonucleotides Coccus-11 (5′–CCA–TCT–TTC–GG–3′) and Coccus(5mC)-11 with methylated cytosines (5′-(5mC)(5mC)A-T(5mC)T-TT(5mC)-GG-3′) were synthesized by Syntol (Moscow, Russia) and were used for the experiments. All oligonucleotides were dissolved in nuclease-free water at a concentration of 100 ng/μL and applied to plant leaves at a rate of 5 mL per m^2^. A control group treated with water was included for comparison. The application rate was 1 mg of DNA in 10 mL of solution per m^2^ of foliage infested with the pest. Oligonucleotide insecticides were directly applied to first- and second-instar larvae. To ensure full coverage, the sprayer angle was adjusted so that the oligonucleotides reached the entire leaf surface harboring the pests. During the treatment, the daytime air temperature was 17 °C, humidity was 84%. Over the two-week experimental period (in November 2025), the average daytime temperature was 16.27 ± 2.11 °C, at night – 13.07 ± 2.41 °C, humidity – 76.94 ± 7.12%. Mortality was calculated by dividing the number of dead individuals by the total number of individuals per leaf and multiplying by 100 to express the result as a percentage. Insects were counted using a Nikon SMZ745 microscope (Nikon Inst., Gyoda, Japan) and images were captured with a Toupcam-UCMOS-05100KPA camera (Asmetec, Kirchheimbolanden, Germany).

### 4.9. Microscopy of leaf surfaces for nucleic acid detection

The studies were performed on temporary anatomical preparations of *P. tobira* (Thunb.) Aiton leaves. For staining nucleic acids, the method of R. Riegler was modified [60] using acridine orange fluorochrome.

Microscopy was performed using a MIKMED-2var.26 fluorescence microscope (LOMO-Microsystems, Saint Petersburg, Russia) and objectives: PLAN F 10x/0.30 and PLAN F 20x/0.50. A block of filters “B” (BP460-490/DM500/BA520) was used, which corresponded to an excitation illumination spectrum of 460-490 nm. A 500 nm barrier filter was used; the spectrum of the recorded fluorescence was 520-700 nm. Photographic recording was performed using a digital camera “MC-6.3” (LOMO-Microsystems, Saint Petersburg, Russia), integrated with the software “ToupView” (ToupTek Photonics, Zhejiang, China).

### 4.10. Differential Gene Expression (DGE) Analysis

RNA quality and quantity were assessed using a BioAnalyser and the RNA 6000 Nano Kit (Agilent Technologies, Santa Clara, CA, USA). PolyA RNA was isolated using the Dynabeads^®^ mRNA Purification Kit (Thermo Fisher Scientific, Waltham, MA, USA). Libraries were prepared using the NEBNext^®^ Ultra™ II RNA Library Prep Kit (New England Biolabs, Beverly, MA, USA). Library concentrations and fragment size were assessed using a Qubit fluorometer (Thermo Fisher Scientific, Waltham, MA, USA) and High-Sensitivity DNA Kit (Agilent Technologies, Santa Clara, CA, USA). Sequencing was performed on an Illumina HiSeq1500 (Illumina, San Diego, CA, USA), generating at least 10 million 50 nt reads per sample. Reads were aligned to the genome using STAR, and differential expression was analyzed using DESeq2 (Bioconductor, Seattle, DC, USA). Reference genome: *Coccus hesperidum*, whole genome shotgun sequencing project (https://www.ncbi.nlm.nih.gov/nuccore/CAXVDF000000000.1) (accessed on 30 March 2026). DGE was performed in two biological replicates for both the Coccus-11-treated and ACTG-11-treated (5′–ACT–GAC–TGA–CT–3′) control groups, with 100 larvae per replicate.

### 4.11. Statistical Analyses

The non-parametric Mann–Whitney U test was used to evaluate the significance of the difference in gene expression. The non-parametric Pearson’s chi-squared test (χ^2^) with Yates’s correction was performed to evaluate the significance of the difference in mortality between control and experimental groups; *p* < 0.01 was considered significant. All calculations were performed using Prism 9 software (GraphPad Software Inc., Boston, USA). All experiments were conducted in triplicate.

## 5. Conclusions

This study demonstrates that the outermost layer of eci-DNA of *P. tobira* leaves is predominantly single-stranded and exhibits a significant insecticidal effect on the pest *C. hesperidum*. Under natural conditions, plant DNA is unwound, partially degraded, and forms a previously unrecognized nanolayer of ssDNA on leaf. This adaptation may serve as a natural defense mechanism against sap-feeding insects. Notably, chloroplast DNA disproportionately enriched in eci-DNA compared to total cellular DNA, suggesting an active release mechanism for chloroplast-derived sequence in eci-DNA formation. The insecticidal activity of natural DNA appears to function through a DNA containment mechanism, wherein eci-DNA interferes with the expression of plant mRNAs and non-coding RNAs (including rRNAs) that share full or partial sequence complementarity. Moreover, oligonucleotide insecticides (DNA insecticides) are capable of activating insect retrotransposons and up-regulating RT-RNase H, crucial enzyme for DNA containment mechanism and causing insecticidal effect. Intriguingly, the ‘genetic zipper’ technology, developed in the laboratory for pest control, has a strikingly similar analogue in nature, underscoring the evolutionary convergence of DNA-based defense strategies. The findings of this study support that CUAD biotechnology is a nature-based pest control solution, reinforcing its safety and potential for agricultural applications. CUADb-based oligonucleotide pesticides hold promise as standalone treatments or as components of integrated bioformulations for the selective management of a wide range of insect pests, ushering in a new era of DNA-programmable plant protection.

## Author Contributions

Conceptualization, V.O.; methodology, V.O., N.G., N.P. and Y.Y.; software, V.O., N.G. and N.P.; validation, V.O. and N.G.; formal analysis, V.O., N.G., N.P. and Y.Y.; investigation, V.O., N.G., N.P. and Y.Y; resources, V.O.; data curation, V.O., N.G., N.P. and Y.Y.; writing—original draft preparation, V.O., K.L., N.G., J.A., N.P., Y.Y. and I.C.; writing—review and editing, V.O., K.L., N.G., J.A., N.P., Y.Y. and I.C.; visualization, V.O., N.G. and N.P.; supervision, V.O..; project administration, V.O.; funding acquisition, V.O. All authors have read and agreed to the published version of the manuscript.

## Funding

The research obtained funding from the Russian Science Foundation No. 25-16-20070, https://rscf.ru/project/25-16-20070/ (accessed on 30 March 2026) (Sections 1, 2, 3, 4 and 5; Sections 2.1, 2.2, 2.3, 2.4, 2.5, 2.7; Sections 4.1, 4.2, 4.3, 4.4, 4.5, 4.6, 4.7, 4.8, 4.10, 4.11), and obtained funding within the framework of a state assignment V.I. Vernadsky Crimean Federal University for 2026 with the planning period of 2024–2026 No. FZEG-2024–0001 (Sections 2, 3, 4 and 5; Section 2.6; Section 4.9).

## Data Availability Statement

The original contributions presented in this study are included in the article/supplementary material. Further inquiries can be directed to the corresponding author.

## Acknowledgments

We thank our many colleagues, too numerous to name, for the technical advances and lively discussions that have prompted us to write this article. We apologize to the many colleagues whose work has not been cited. We are very much indebted to all anonymous reviewers and our colleagues from the Lab on DNA technologies, PCR analysis and creation of DNA insecticides (V.I. Vernadsky Crimean Federal University, Institute of Biochemical Technologies, Ecology and Pharmacy, Department of General Biology and Genetics), and OLINSCIDE BIOTECH LLC. for valuable comments on our manuscript.

## Conflicts of Interest

The authors declare no conflicts of interest.

## Disclaimer/Publisher’s Note

The statements, opinions and data contained in all publications are solely those of the individual author(s) and contributor(s) and not of MDPI and/or the editor(s). MDPI and/or the editor(s) disclaim responsibility for any injury to people or property resulting from any ideas, methods, instructions or products referred to in the content.

## Notes

### Competing Interest Statement

The authors have declared no competing interest.

### Summary of Updates

Revised Title; Revised Section 2; added subsection 2.7 and 4.10; added Table 1; updated Figure 10; Added Graphical abstract. Reference revised and updated.

